# Dual RNA-Seq analysis of *in vitro* infection multiplicity in *Chlamydia*-infected epithelial cells

**DOI:** 10.1101/2020.10.21.347906

**Authors:** Regan J. Hayward, Michael S. Humphrys, Wilhelmina M. Huston, Garry S.A. Myers

## Abstract

Dual RNA-seq experiments examining viral and bacterial pathogens are increasing, but vary considerably in their experimental designs, such as infection rates and RNA depletion methods. Here, we have applied dual RNA-seq to *Chlamydia trachomatis* infected epithelial cells to examine transcriptomic responses from both organisms. We compared two time points post infection (1 and 24 hours), three multiplicity of infection (MOI) ratios (0.1, 1 and 10) and two RNA depletion methods (rRNA and polyA). Capture of bacterial-specific RNA were greatest when combining rRNA and polyA depletion, and when using a higher MOI. However, under these conditions, host RNA capture was negatively impacted. Although it is tempting to use high infection rates, the implications on host cell survival, the potential reduced length of infection cycles and real world applicability should be considered. This data highlights the delicate nature of balancing host-pathogen RNA capture and will assist future transcriptomic-based studies to achieve more specific and relevant infection-related biological insights.

## Introduction

Dual species transcriptomic experiments (dual RNA-seq) allow multiple organisms to be simultaneously analysed from within the same sample, such as host and bacterial transcripts during an infection [1]. The number of dual RNA-seq experiments are increasing, with the underlying experimental designs and infection conditions often varying significantly, particularly in infection rates and RNA depletion methods [2-7].

Here, we have applied dual RNA-seq to human epithelial cells subjected to the bacteria *Chlamydia trachomatis*, which is an obligate intracellular, human-specific bacterial pathogen that causes trachoma and urogenital infections [8-10]. Ocular infections cause trachoma (infectious blindness), typically in disadvantaged communities, and is the leading cause of preventable blindness worldwide [9, 10]; while genital infections are the most prevalent sexually transmitted infection (STI) worldwide [9]. If infections are left untreated they can become problematic leading to more complex disease outcomes including ectopic pregnancy and infertility [11, 12]. Diagnosed chlamydial infections can successfully be treated with antibiotic therapy, however asymptomatic infections are common [13, 14] and thus challenging to treat. Genome-wide transcriptomic studies have explored gene expression from infected host cells and chlamydial-specific expression, either separately or simultaneously [15-23]. The previous chlamydial-based dual RNA-seq experiment encompassed an experimental design that used a multiplicity of infection (MOI) of 1, while their depletion technique removed rRNAs in all samples, followed by subjecting half of these libraries to polyA depletion to further enrich chlamydial transcripts. Although two depletion methods were used, it is uncertain if this increased the abundance of chlamydial transcripts. Additionally, an MOI of 1 at an early time point highlighted low capture rates of chlamydial transcripts [15].

In general, host-based RNA-seq experiments in an infection setting will typically try and achieve a ratio of 1 infectious entity per host cell. This ratio is referred to as the MOI, with an MOI of 1 indicating a 1:1 ratio, and is frequently used to assess baseline changes in both organisms without any directional bias. RNA-seq and microarray experiments that have focused on chlamydial infection have utilised a range of MOIs ranging from 1 [24] to 100 [16, 20]; with higher ratios helping to exaggerate and highlight the chlamydial impact. However, too high an MOI and the whole monolayer of cells dies before the infection can proceed. In addition, a higher MOI will likely affect the developmental cycle due to the underlying stress this places on host cells [25, 26].

In this experiment, both host and chlamydial gene expression were examined applying dual-RNA-seq to *in vitro C. trachomatis*-infected HEp-2 epithelial cells. The first aim was to understand the influence different MOIs have on sequence capture rates, but also the transcriptional variation from *Chlamydia* and the host-cell. The second aim attempted to improve the enrichment of chlamydial reads by comparing different RNA depletion methods. To address these questions, two time points were chosen covering the chlamydial developmental cycle (1 and 24 hours), with each time point split into three MOIs (0.1, 1 and 10), each in triplicate. Each of these biological replicates (16 samples) were split in half, where one library was prepared solely with rRNA depletion, while the second was prepared with rRNA depletion followed by polyA depletion (**Figure 1A**).

**Figure 1:**
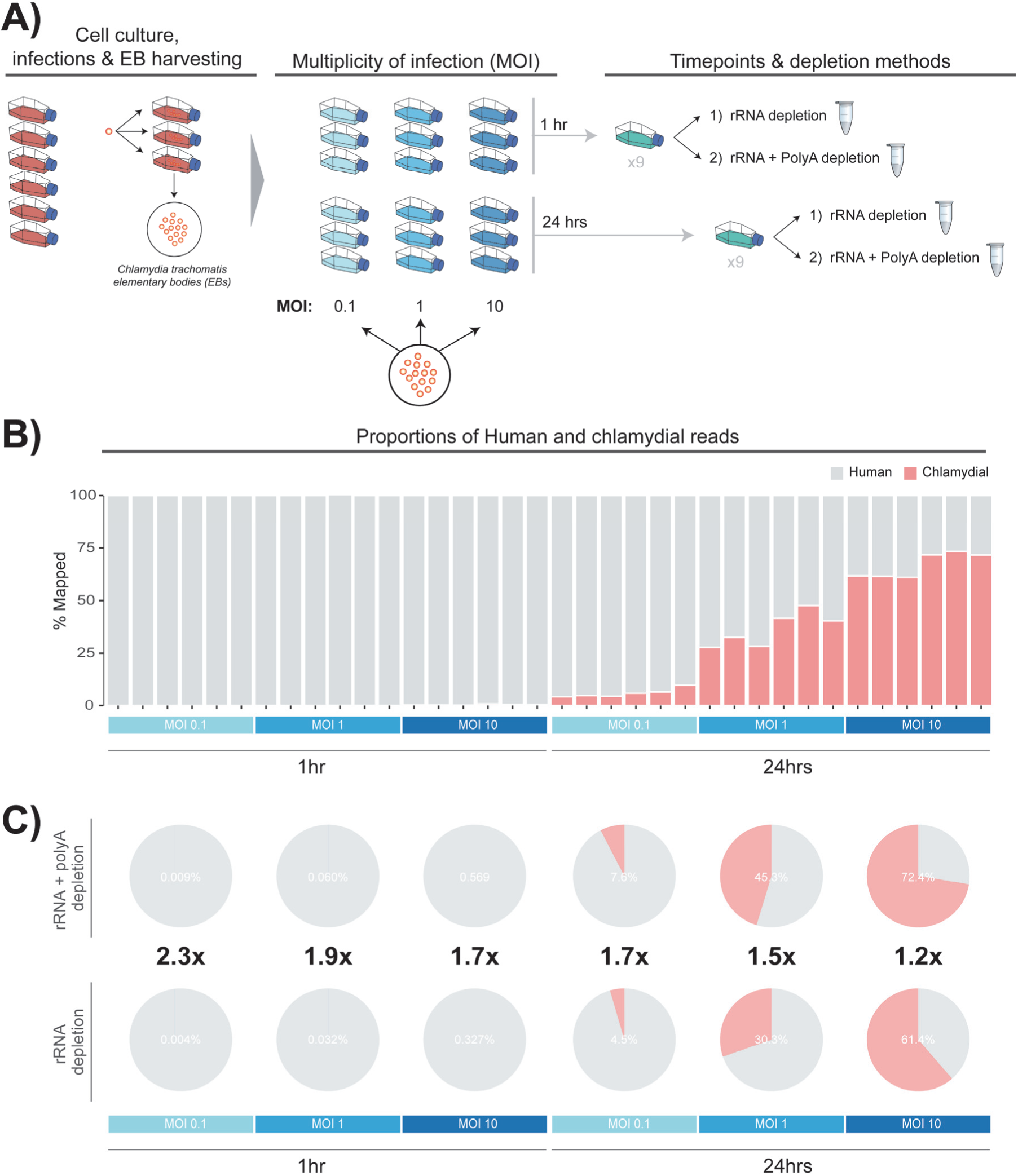
Experimental process and design. **A)** The process of growing cell cultures and harvesting (elementary bodies) EBs to use for downstream experiments is a time-consuming process spanning multiple days. The resulting EBs were used for three different infection ratios (multiplicity of infection) of 0.1, 1 and 10. After infections, samples were left for 1 hour and 24 hours. Each replicate was then prepared with rRNA or rRNA plus PolyA depletion, generating 32 samples in total. **B)** Showing the percent of Human and chlamydial reads across the experimental design. **C)** By combining rRNA depletion and polyA depletion, we were able to increase the capture efficiency of chlamydial transcripts at both time points and across MOIs

## Methods

### Cell culture and infection

Human epithelial type 2 (HEp-2) cells (American Type Culture Collection, ATCC No. CCL-23) were grown as monolayers in 6x 100 mm tissue culture dishes until cells were 90% confluent. To harvest EBs for the subsequent infections, additional monolayers were grown and infected with *C. trachomatis* serovar E in sucrose phosphate glutamate (SPG) as previously outlined [27]. The resulting EBs and cell lysates were then harvested and used to infect new HEp2 monolayers.

Infections for each dataset used the previously prepared HEp2 monolayers, infecting with *C. trachomatis* serovar E in 3.5 mL SPG buffer as previously outlined [27]; infections were synchronised using centrifugation. EBs were introduced into monolayers from three MOIs (0.1, 1 and 10) using 1:10 dilutions beginning from an MOI of 10. EBs were quantified as previously described [15]. To remove non-viable or dead EBs, each sample was incubated at 25°C for 2 hours, and washed twice in SPG. Cell monolayers were incubated at 37°C with 5% CO2, including the addition of 10 mL fresh medium (DMEM+10% FBS, 25 μg/ml gentamycin, 1.25 μg/ml Fungizone). After each infection time point, the infected and uninfected dishes were harvested by scraping and resuspending in 150 μL sterile PBS. Any resuspended samples were stored at −80°C.

### Library preparation and sequencing

Ribo-Zero rRNA Removal kits (Human/Mouse/Rat and Gram-negative) were used to deplete samples of both human and gram-negative bacterial rRNA. Equivalent volumes from each kit were combined, thereby allowing the removal of bacterial and human rRNA simultaneously within each sample. Each sample was equally separated, with one half subjected to polyA depletion by the Poly(A) Purist Mag purification kit (Ambion), whereby removing host-based polyA transcripts to allow the enrichment of bacterial transcripts. Magnetic beads were used to bind to polyA mRNAs and were extracted from the solution with a magnet. Samples with combined depletion methods were further purified using Zymo-Spin IC columns (Zymo Research) before being re-combined for library construction.

The mRNA libraries were prepared from depleted samples as previously stated at 1 and 24 hours post infection, using the TruSeq RNA Sample Prep kit (Illumina, San Diego, CA) per the manufacturer’s protocol with IGS-specific optimisations. Adapters and indexes (6 bp) were ligated to the double-stranded cDNA, which was subsequently purified with AMPure XT beads (Beckman Coulter Genomics, Danvers, MA) between enzymatic reactions and size selection steps (∼250 to 300 bp). The resulting libraries were sequenced on an Illumina HiSeq2000 using the 100 bp paired-end protocol at the Genome Resource Centre, Institute for Genome Sciences, University of Maryland School of Medicine.

### Bioinformatic analysis

Sequencing reads were trimmed and quality checked using Trim Galore (0.45) (https://www.bioinformatics.babraham.ac.uk/projects/trim_galore/) and FastQC (0.11.5) [28]. Host reads were aligned to the human genome (GRCh 38.87) using STAR (2.5.2b) [29], while chlamydial reads were aligned to the *Chlamydia trachomatis* (serovar E, Charm001) genome using Bowtie2 (2.3.2) [30] with additional parameters of ‘1 mismatch’ and ‘–very-sensitive-local’. Samtools (1.6) [31] was used to remove duplicate reads in addition to only keeping mapped reads in both the host and chlamydial BAM files. To remove reads that mapped to both genomes, we first extracted the mapped reads back into paired-end fastq files using bedtools (2.26.0) [32]. Reads were then aligned using the initial mapping software to the reciprocal genomes. Any reads that mapped to both genomes were removed from the originating BAM files using the “FilterSamReads” command from Picard tools (2.10.4) [33]. Additional quality control metrics were examined using Bamtools (2.5.1) [34], MultiQC (1.2) [35] and various in-house scripts.

Features (genes) were counted using featureCounts (1.5.0-p1) [36] with additional parameters of “-Q 10 -p -C”. Genefilter (1.64.0) [37] was used to filter out genes with low counts, where host genes were retained if expression > 50 in at least three samples. To accommodate the vast differences in expression between host and chlamydial reads, a separate filter was used retaining chlamydial genes with expression > 10 in at least three samples. Chlamydial and host reads were further separated by time point due to the large amount of variability in expression between an MOI of 0.1 at 1 hour to an MOI of 10 at 24 hours. Once separated, library normalisation was performed using the trimmed mean of M-values (TMM) method [38].

To identify outliers, four PCA bi-plots were generated from library normalised counts using PCATools (0.99.13) [39], where eigenvalues from PC1 and PC2 for each replicate were calculated and used to highlight outlier samples if an eigenvalue was > |3| standard deviations from the mean within that group. If an outlier was removed, eigenvalues were recalculated and the process repeated until no further outliers were detected. To determine the underlying variation at each principal component, the “plotloadings” function within PCATools [39] was used.

Differential expression was performed with edgeR (3.24.3) [40], adding the difference between the depletion methods as a blocking factor, whereby allowing MOI and time point comparisons to utilise all six replicates to increase significance. Host DE genes were uploaded and enriched for KEGG pathways using the Enrichr database [41]. Relevant host pathways were determined using the combined score with a cutoff of > 50. Combined scores were calculated by adding together the combined scores comparing MOIs 0.1 vs 1, and 1 vs 10. Enrichment of chlamydial DE genes was performed using the ‘UniProt Keywords’ feature of STRING (11.0) [42].

### Data availability

The data set supporting the results of this article is available in the GEO repository, GSE150039.

## Results

### Quantifying expression differences between host and chlamydial reads

Dual RNA-seq was applied to *C. trachomatis* serovar E-infected human HEp-2 epithelial cells in triplicate at 1 and 24 hours post-infection (hpi). Within each time point, three MOIs were used (0.1, 1 and 10), in addition to two depletion methods 1) rRNA depletion, 2) rRNA depletion and polyA depletion; totalling 36 samples across the experimental design.

Capture rates of chlamydial reads at early time points is challenging due to limited biological activity, where the majority of transcripts in a sample (>99%) will be associated with the host [15]. We increased the sequencing depth at 1 hour (> 6 fold) to try and capture more chlamydial reads; generating 391,847,337 mapped reads at 1 hour compared to 63,710,236 mapped reads at 24 hours. Even with this greater depth of sequencing at 1 hour, the number of chlamydial reads was still quite low; especially at an MOI of 0.1 with an average of 1,407 reads across the six replicates. However, as the MOI increases, we do see an increase in chlamydial transcripts, with average mapped reads of 10,392 (MOI 1), and 55,426 (MOI 10) (**Figure 2A**). At 24 hours we see an expected increase in the number of chlamydial reads, following a similar trend with 1 hour, where the number of mapped reads increases as the MOI increases (**Figure 2B**). The number of mapped host reads tends to vary more than the chlamydial reads, particularly between depletion methods of the same MOI (**Figure 2C** and **D**). This is likely due to the increased variety of host transcripts resulting from post-transcriptional modifications, such as polyadenylation, which does not occur in bacterial systems.

**Figure 2:**
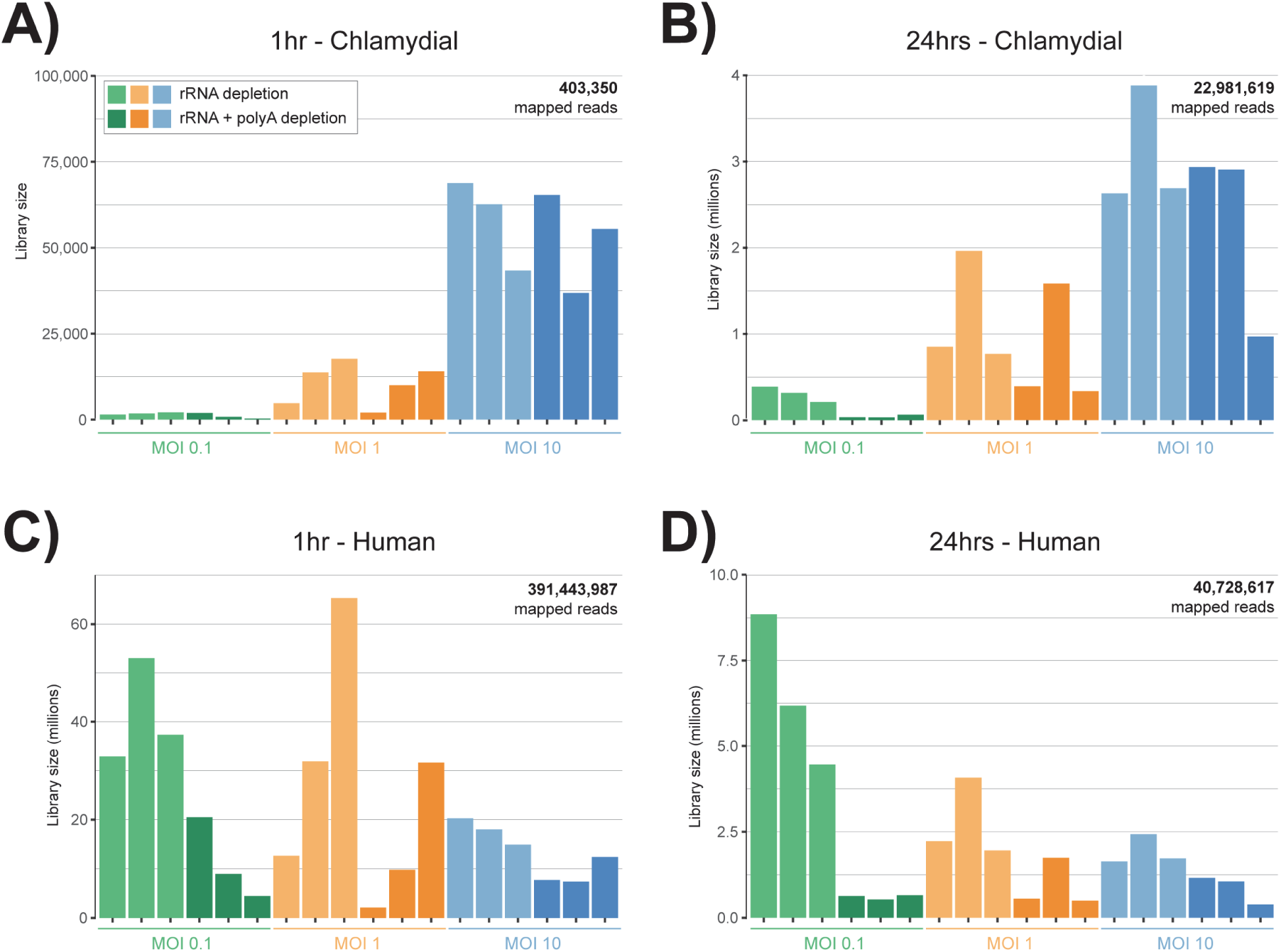
Human and chlamydial mapped reads. The number of mapped sequence reads to both human and chlamydial genomes. Green bars represent an MOI of 0.1, orange 1, and blue 10. Light shaded colours represent the rRNA depletion method, while darker shades represent rRNA and polyA depletion methods combined. **A)** Low numbers of chlamydial mapped reads at lower MOI ratios, but a substantial increase at an MOI of 10. **B)** At 24 hours the number of captured transcripts dramatically increases from 1 hour, but follows a similar distribution with a spike of reads at an MOI of 10. **C)** The increase in mapped host reads at 1 hour was to try and capture as many chlamydial transcripts as possible, as they are known to be in very low quantities this early in the developmental cycle. **D)** As the MOI increases at 24 hours, the number of mapped host reads declines.

When examining the proportions of host and chlamydial reads together across the experimental design, we see that 1 hour is dominated by host reads, while at 24 hours we see a gradual increase of chlamydial reads as the MOI increases. Surprisingly, at 24 hours with an MOI of 10, the proportion of chlamydial reads across all replicates is over 60% (**Figure 1B**).

### Combining depletion methods increases yield of bacterial transcripts

By combining two depletion methods (rRNA depletion and polyA depletion), we had anticipated capturing additional chlamydial reads. The addition of polyA depletion should theoretically remove any polyadenylated host transcripts, thereby increasing the number of chlamydial transcripts to be captured and sequenced.

Overall, we see an increase in chlamydial reads when combining depletion methods. Even at 1 hour when there are limited transcripts circulating within the cell, we still see an average increase of 2.0x. At 24 hours when more chlamydial transcripts are being expressed, we see an average increase of 1.5x more reads. Interestingly, at 24 hours as the MOI increases, the capture efficiency begins to decline slightly from 1.7x to 1.2x (**Figure 1C**).

### Differences in chlamydial expression between depletion methods

PCA bi-plots were created to compare the expression profiles across replicates from both depletion methods. At 1 hour, we see minimal separation at an MOI of 1 and 10 compared to 0.1 where replicates appear separated and not grouped by depletion method as expected (**Figure 3A**). However, none of the replicates were considered outliers using a robust statistical approach as outlined in the methods. We therefore attribute this variability to the low number of chlamydial reads present at an MOI of 0.1 as identified earlier. At 24 hours a distinct separation between depletion methods within each MOI can be easily visualised (**Figure 3B**).

**Figure 3:**
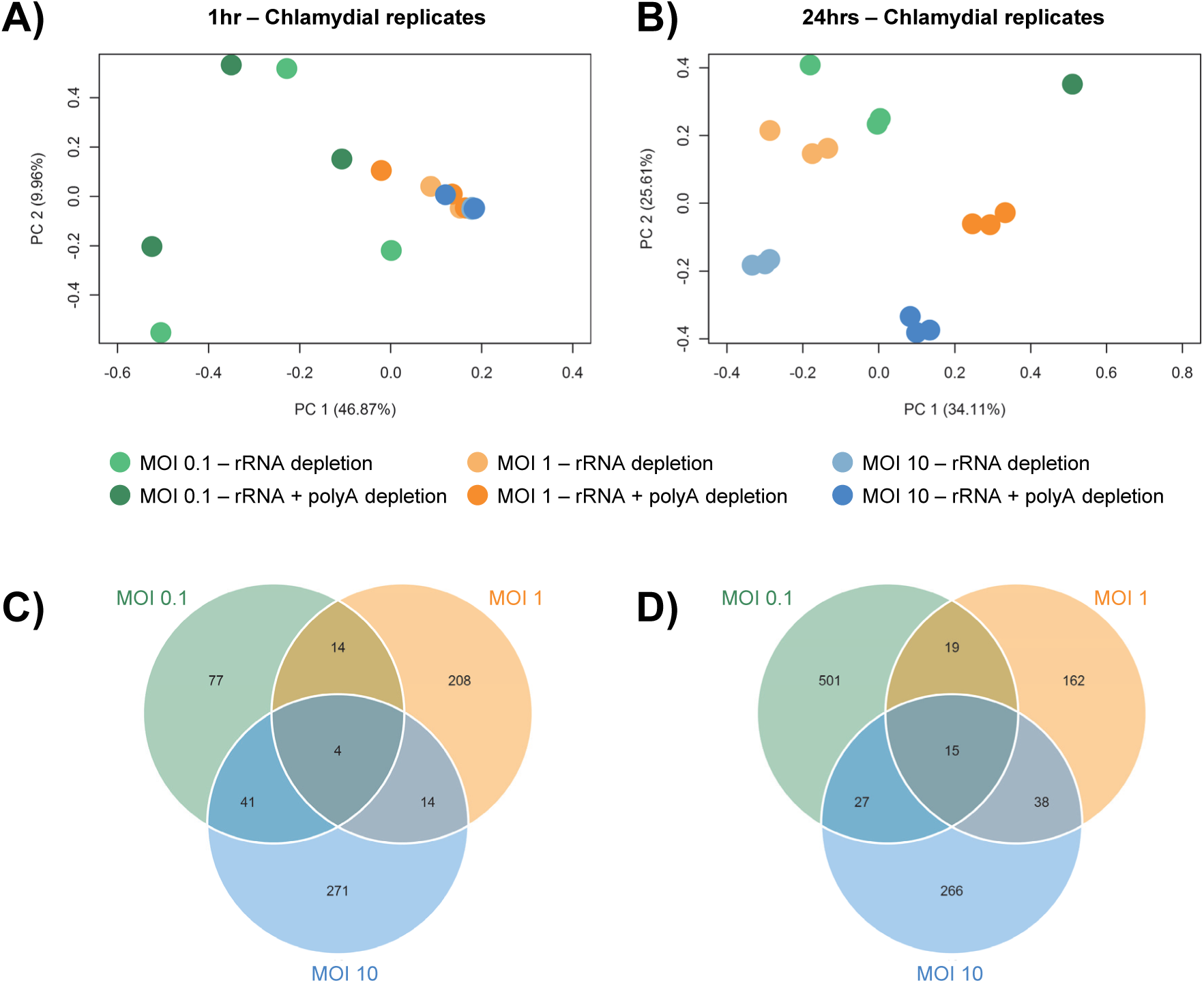
Chlamydial-based expression differences between depletion methods. **A)** At 1 hour, minimal separation is seen apart from at an MOI of 0.1 where replicates appear to not group or cluster together. **B)** At 24 hours, replicates group together within the same depletion method, while separation is seen between depletion methods at each MOI. Extracting the top 5% of genes driving variation from PC1 and PC2 between depletion methods. At 1 hour **C)** and 24 hours **D)**, we see subsets of genes overlapping MOIs in addition to MOI-specific subsets.

To understand if the variability between depletion methods is driven by a small subset of highly expressed genes, or an assortment of genes, we extracted the top 5% of genes driving the underlying variation at PC1 and PC2 for each MOI (**Figure 3C-D**). At both time points we see subsets of genes specific to each MOI, indicating that each MOI exhibits a slightly different chlamydial response. In addition, overlapping genes highlight that the variation between depletion methods was also captured and overlaps considerably. Therefore, the inclusion of polyA depletion increases bacterial reads and does not seem to be driven by small subsets of highly expressed transcripts, but allows for a wide array of transcripts to be captured.

### The removal of polyA transcripts increases non-protein coding host gene expression

Examining PCA bi-plots for host reads show tight clustering between replicates, but also highlights the separation between depletion methods (**Figure 4A-B**). Extracting the underlying genes contributing the variation at PC1 and PC2, numerous non-coding genes were identified. To calculate the percent of protein coding versus non-protein coding expression, gene expression was averaged across replicates after separation by time point, MOI and depletion method (**Figure 4C**). Across both time points we see an average of 2.8x more non-protein coding expression when combining rRNA and polyA depletion, with the highest proportion occurring at an MOI of 0.1 (3.4x at 1 hour and 4.9x at 24 hours) (**Figure 4C**). Pie charts highlight that the majority of expression comes from protein-coding genes. However, as identified in the PCA plots earlier, non-protein coding expression contributed to the separation of depletion methods. By characterising the most common biotypes, we see mitochondrial rRNA (MT rRNA), small nucleolar RNAs (snoRNA), miscRNA and long intergenic non-coding (lincRNA); but without any visible trends separating time points, depletion methods or MOI (**Figure 4D**).

**Figure 4:**
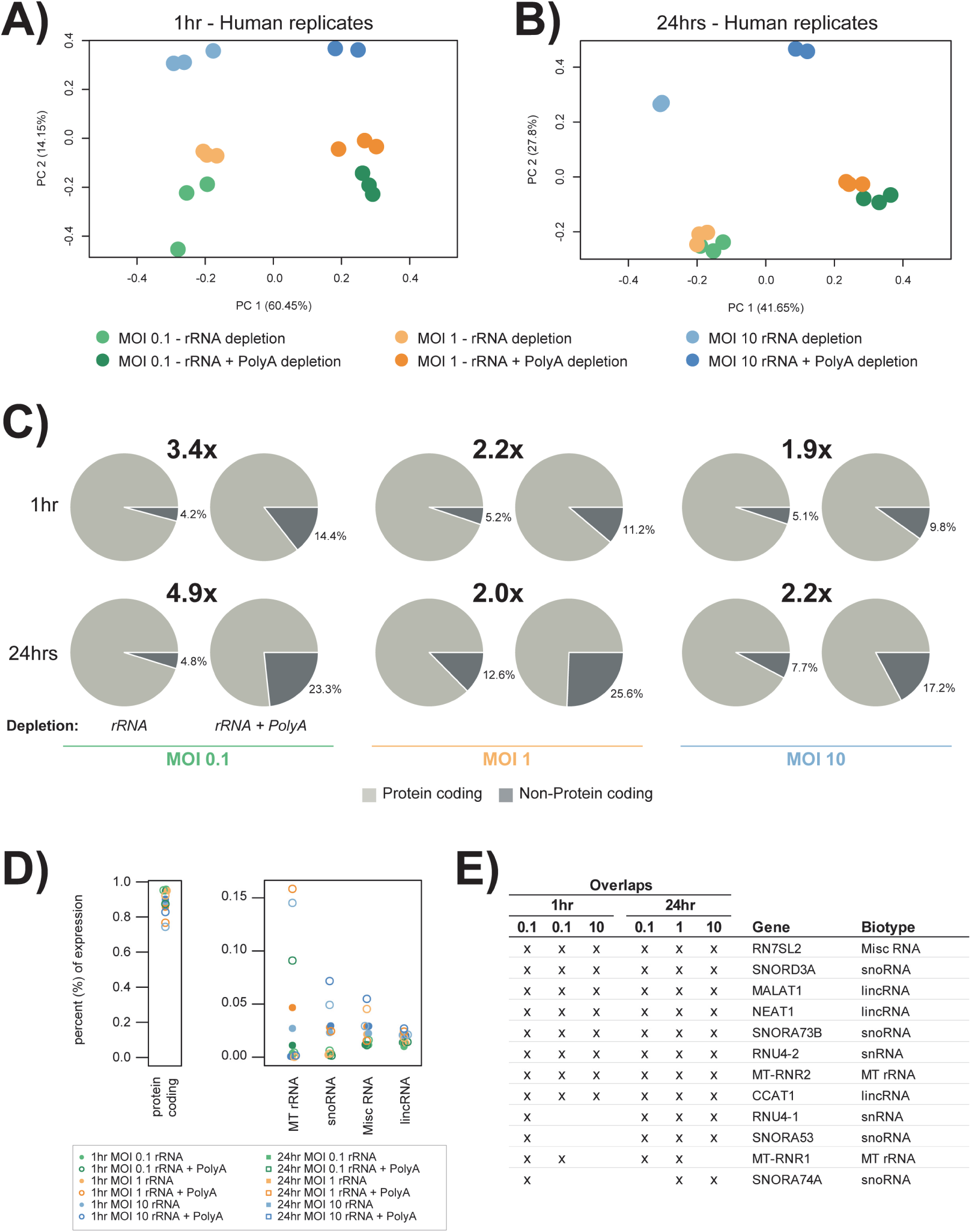
Host-based expression differences of protein coding and non-protein coding genes between depletion methods. **A)** PCA plots show tight grouping between replicates, but separation at each MOI and depletion method. **B)** Similar grouping trends to 1 hour, but with an MOI of 10 much further separated. **C)** An overall increase in non-protein coding expression is observed when combining depletion methods. **D)** The majority of expression for all conditions is from protein coding genes. While non-protein coding genes are dominated by four biotypes, with Mt rRNAs the most highly expressed. **E)** Non-protein coding genes that are within the top 200 expressed genes at each MOI, and overlap 3 or more conditions.

To identify potentially influential non-protein coding genes, we used the top 200 expressed genes from both depletion methods and extracted a subset of genes that occur frequently (across 3 or more conditions) (**Figure 4E**). Of the 12 genes identified, 5 were snoRNAs which are involved with RNA modifications, and are among the most highly abundant non-coding RNAs (ncRNAs) in the nucleus [43]. The MT-RNR1 (12S RNA) and MT-RNR2 (16S RNA) genes encode the two rRNA subunits of mitochondrial ribosomes, and are generally always highly expressed within eukaryotic cells [44]. LincRNAs include CCAT1, which is linked to cell growth and regulation of EGFR [45], while MALAT1 and NEAT1 co-localise to hundreds of genomic loci, predominantly over active genes [46].

### Increasing infection highlights minimal changes to highly expressed host and chlamydial genes

To determine whether the host or chlamydial transcriptional-profile changes in relation to the ratio of EBs per cell, highly expressed genes were compared against an MOI of 1. Chlamydial transcripts were examined from the combined depletion replicates, as more transcripts were captured (**Figure 1C**), thus giving a more representative profile. Host reads were taken from just the rRNA depleted replicates, as these were shown to contain more of an accurate representation of protein coding and non-protein coding genes (**Figure 4**).

Each of the four panels (**Figure 5**) contains two graphs. The first graph contains the top 50 expressed genes taken from an MOI of 1, while the second graph shows the ranked-positions of these top expressed genes. At 1 hour, there is slightly less chlamydial expression at an MOI of 0.1, and slightly more expression when additional EBs are introduced at an MOI of 10 (**Figure 5A**). The ranking chart to the right shows that 9/10 of the top expressed genes remain the same across the three MOIs. The top 25 genes from the host’s response at 1 hour share highly similar expression profiles (**Figure 5B**); with only two mitochondrial-based genes (MT-RNR1 and MT-RNR2) at an MOI of 10 standing out with lower expression. The ranking chart shows 7/10 top expressed genes remaining constant across the three MOIs, similar to the chlamydial profile. At 24 hours, similar expression profiles of the top 25 expressed chlamydial genes are seen, irrespective of MOI (**Figure 5C**). Rankings are also similar, with only slight variations in the top ten genes, and 16/20 of the top expressed genes remain identical. The host expression profile at 24 hours is consistent at an MOI of 1 and 0.1, whereas the expression pattern is more widely distributed at an MOI of 10; again with MT-RNR1 and MT-RNR2 exhibiting lower expression (**Figure 5D**). Although the top ranked genes exhibit more variability within their rankings compared to 1 hour, 90% of the top expressed host genes appear at both time points. Functional characterisation of the genes shows their involvement with general cell-based growth events, such as ribosomal-based processes, metabolism, and cytoskeletal components (**Supplementary File 1**). Many top ranked host genes are also non-protein coding as identified by an asterisk (*). However, with limited annotation available, their characterisation in to infection-association functions are limited. Of the annotated non-protein coding genes, they appear to be involved with general cell regulatory processes. Only seven chlamydial genes overlap both time points, which was anticipated, as two different biological events are occurring at these times, including infection mechanisms at 1 hour, and growth-related processes at 24 hours. Characterisation of these overlapping genes identifies membrane proteins and transcription/translation machinery, which are needed throughout the developmental cycle (**Supplementary File 1**).

**Figure 5:**
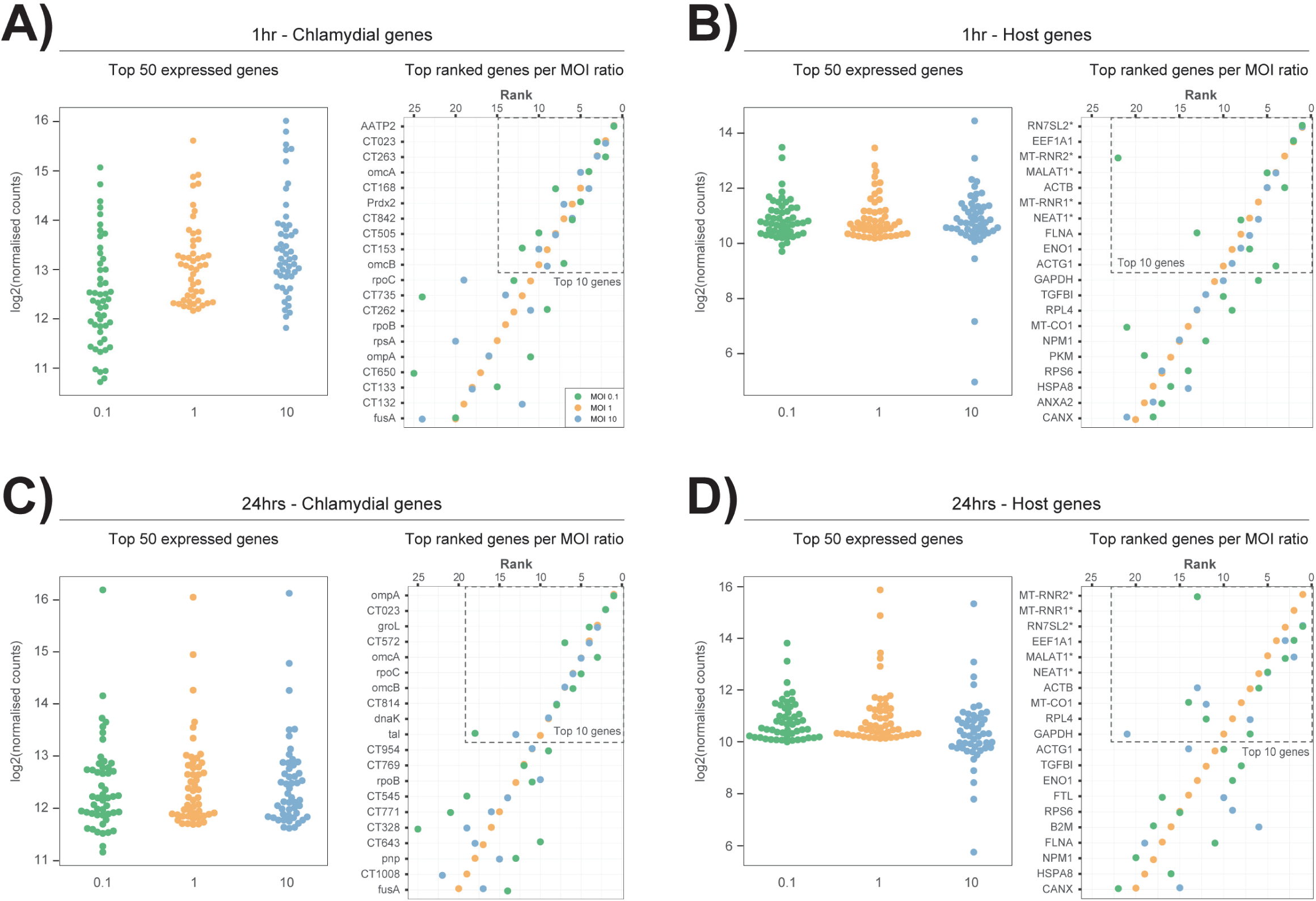
Top 25 expressed host and chlamydial genes across MOIs. Each quadrant contains two graphs: the first showing the top 50 expressed genes across each MOI (taken from an MOI of 1), while the second ranks those same genes for direct comparison. **A)** Chlamydial expression at 1 hour shows a slight upwards trend as the MOI increases, with high similarity in the ranked order. **B)** Host expression at 1 hour shows consistent expression across MOIs apart from two mitochondrial genes (MT-RNR1 and MT-RNR2) with low expression at an MOI of 10. Rankings of the top ten genes are also highly similar. **C)** At 24 hours, chlamydial expression seems to be less influenced by MOI. **D)** Host expression at 24 hours shows a slight increase in overall expression, while the MOI of 10 has begun to have more of a varied range of expression compared to 1 hour. Ranked genes also remain highly similar.

### Comparative analysis between MOIs show increased expression of inflammatory and immune-based host genes

To observe and compare how the host expression responded to increased infection, we examined differentially expressed (DE) genes comparing MOIs. At 1 hour, the majority of genes (87% from 0.1 to 1, and 67% from 1 to 10) exhibited an increase in regulation as the MOI increased (**Figure 6A**). By enriching DE genes which are up-regulated and overlap both comparisons, pathways that exhibit an increase in expression as the MOIs increase were identified (**Figure 6B**). The same method was applied to down-regulated genes. However, no continuously down-regulated pathways were identified. The top four up-regulated pathways highlight similar host immune regulated functions that include *(TNF signalling), (NF-κB signalling), (NOD-like receptor signalling)* and *(Cytokine-cytokine receptor interaction)*; with the proinflammatory cytokine TNF exhibiting almost double the combined score of the next highest. We had anticipated seeing a strong immune-based response, as these pathways are associated with primary defence mechanisms; so, they should increase as the bacterial threat increases.

**Figure 6:**
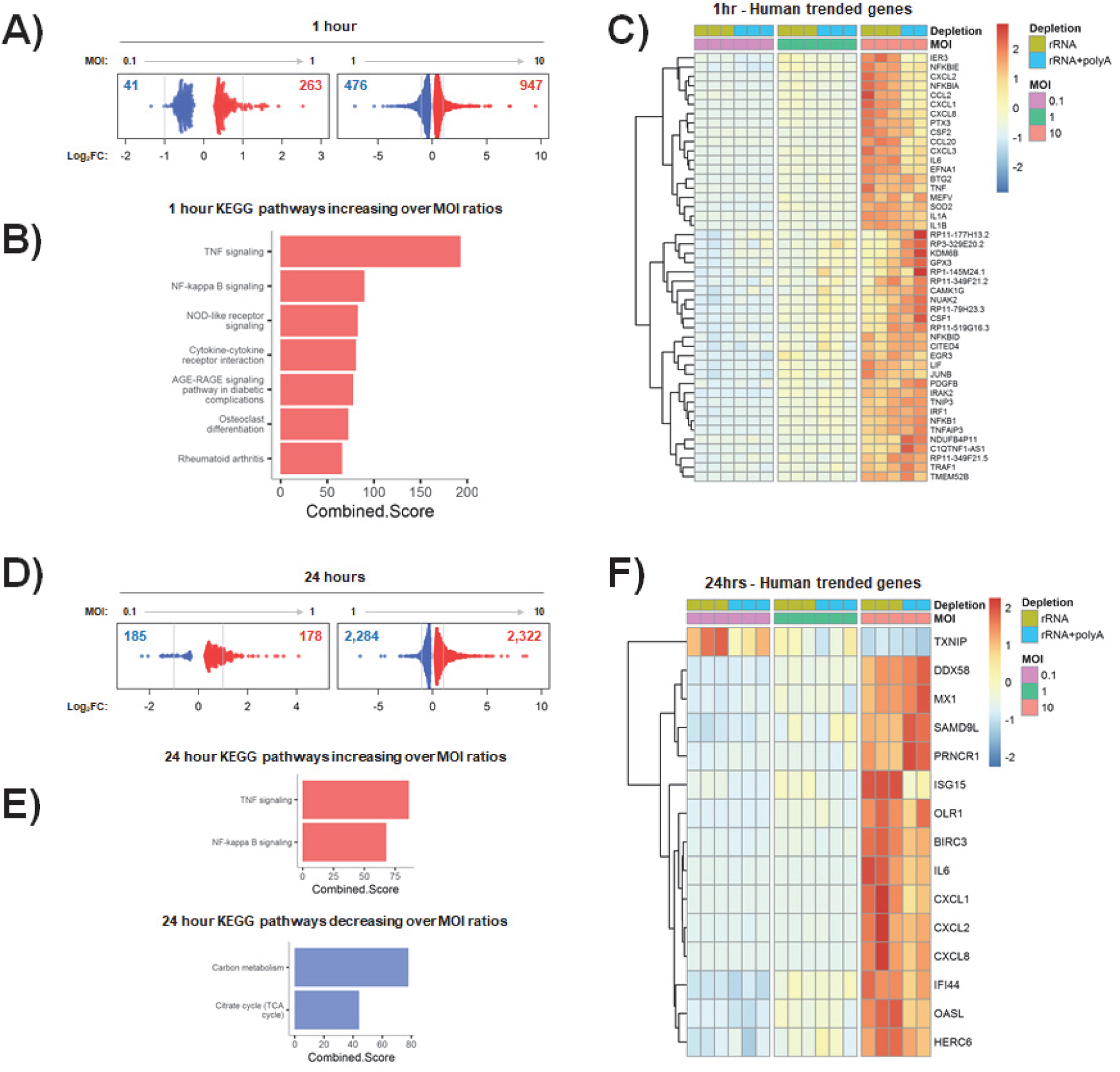
Comparison of differentially expressed host genes across MOIs. **A)** An increase of up-regulated genes at both comparisons is seen at 1 hour. **B)** Extracting and enriching genes that overlap both comparisons and are also up-regulated, show pathways involved with immune and inflammatory responses. **C)** Trended genes are determined from exhibiting a fold-change > 2 and following the same regulation pattern at both comparisons. At 1 hour, 46 up-regulated ‘trended-genes’ further highlight the association with the immune system with over 50% of genes grouped in to cytokine signalling, chemokines and interleukins. **D)** At 24 hours the numbers of up and down-regulated genes is much more even (49% and 50%) than 1 hour. **E)** The top two up-regulated pathways are repeated at both time points, while down-regulated pathways are associated with metabolism and likely indicate cells shifting into defence mode as infection increases. **F)** Trended-genes further highlight immune responses, in addition to viral and ubiquitin-related immune responses.

To further examine influential genes underlying these pathways, ‘trended-genes’ were extracted. The criteria consisted of an expression profile that at least doubled (fold-change >2) for each comparison, in addition to showing a continued increase from an MOI of 0.1 to 10. In total, 46 genes were identified that trended-upwards (**Figure 6C**), no genes trended downwards. These trended-genes further highlight that the underlying host-mechanisms to increased infection at initial stages are predominately immune system associated 24/46 (52%), encompassing cytokine signalling, chemokines and interleukins.

The number of DE genes at 24 hours show an even distribution of fold-changes compared to 1 hour, with 49% up-regulated comparing MOIs 0.1 and 1, and 50% comparing 1 and 10 (**Figure 6D**). Enriched pathways that are continuously up-regulated include *(TNF signalling)* and *(NF-κB signalling)*, which are the same top two pathways found at 1 hour, and strongly linked to inflammation [47]. We also see two enriched pathways that become down-regulated as the MOI increases: *(Carbon metabolism)* and the *(Citrate cycle (TCA cycle))* (**Figure 6E**). This decrease in key metabolism is likely due to cells prioritising defence over growth as the infection escalates.

Examining trended-genes at 24 hours uncovers 1 gene exhibiting decreased expression (TXNIP), and 14 genes with increased expression (**Figure 6E**). TXNIP (Thioredoxin Interacting Protein) is a thiol-oxidoreductase involved in redox regulation which protects cells against oxidative stress [48]. Chlamydial-specific studies have identified an increase in reactive oxygen species (ROS) at early time points, but expression is rapidly reduced shortly afterwards [49]. A further study has suggested that the redox state within a cell could be a regulator in *Chlamydia*-induced apoptosis [50]. However, it is difficult to know if this decreased regulation is directly linked to chlamydial infection and what advantages an oxidative cellular environment would provide at this developmental stage. Genes with increased expression fall into three main categories: cytokines and inflammation (6 genes), viral-based immune response (5 genes), and ubiquitin-related immune responses (3 genes). As anticipated and seen at 1 hour, expression of key immune related genes increases with an increased burden. Only 4 genes overlapped both time points that also increased expression across MOIs (CXCL1, CXCL2, CXCL8 and IL6), indicating their importance as immune mediators against infection.

### Comparative analysis of chlamydial expression between MOIs

DE genes were also identified to explore chlamydial-based changes attributed to different MOIs. The number of DE genes at 1 hour reflected the underlying minimal expression profiles already identified (**Figure 2**), with 47 DE genes comparing MOIs 0.1 and 1, and 23 genes comparing 1 and 10 (**Figure 7A**). At 24 hours, the increase in underlying expression resulted in an increase in DE genes, with 81 comparing MOIs 0.1 and 1, while over half (56%) of the chlamydial genome (566/1008 genes) showed a significant change in regulation comparing MOIs 1 and 10 (**Figure 7B**).

**Figure 7:**
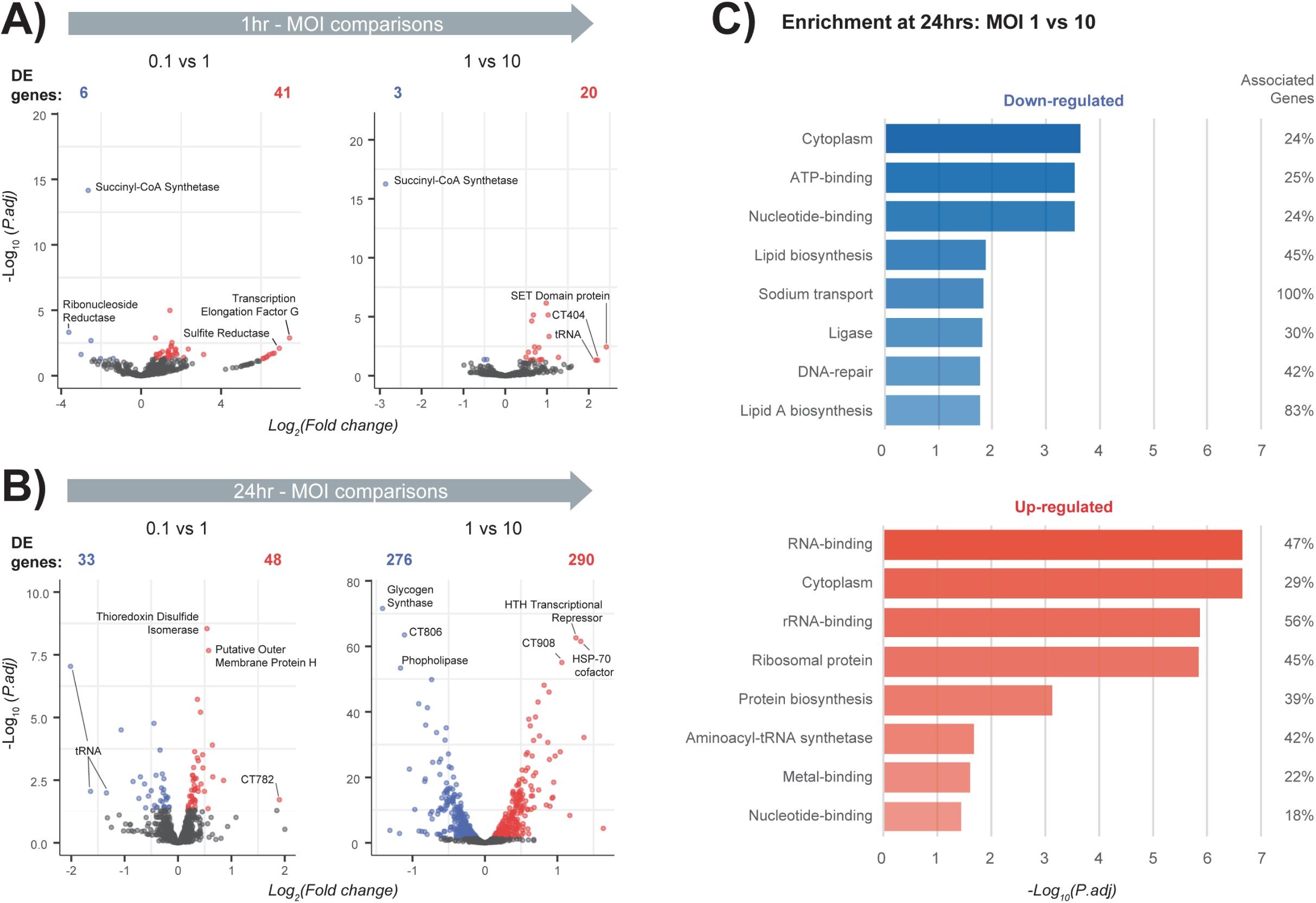
Comparison of differentially expressed chlamydial genes across MOIs. **A)** Volcano plots after differential comparisons between MOIs at 1 hour show minimal differentially expressed (DE) genes. **B)** Differential comparisons at 24 hours show a slight increase in DE genes comparing MOIs 0.1 and 1, with a further considerable increase comparing ratios 1 and 10. **C)** The increase in DE genes comparing MOIs 1 and 10 at 24 hours allowed enrichment of up and down-regulated genes.

No chlamydial genes continuously increased across MOIs at either time point. Only two genes showed a continued decreased in expression: SCLA1|TEF25 (Succinyl-CoA Synthetase) at 1 hour, and CT726 (tRNA) at 24 hours. The decrease of transfer RNAs (tRNA) at 24 hours is slightly surprising, considering they are an important component of translation, and would likely be in abundance during this growth phase of the developmental cycle. Also surprising is a decrease in Succinyl-CoA synthetase, which is involved with the citric acid cycle and cellular metabolism [51]. We can theorise from these genes, as more EBs are introduced, the likelihood of multiple infections within a cell is greatly increased and perhaps some inclusions are benefitting from effector proteins already circulating within the cell from existing inclusions.

Due to low numbers of DE genes at 3 of the 4 comparisons, enrichment was only possible comparing MOIs 1 and 10 at 24 hours (**Figure 7C**). Down-regulated functions comprise genes that show decreased expression at a higher MOI. Results also show unexpected functions such as ‘*ATP-binding*’ and ‘*Lipid biosynthesis*’, which would generally be associated with chlamydial growth. This may highlight the possibility that inclusions may benefit from effector proteins already in existence, likely reducing the need to express these genes and associated processes. Up-regulated genes cover a wider range functions, with half associated with different binding mechanisms facilitating transcription and growth (*RNA-binding, rRNA-binding, Metal-binding* and *Nucleotide-binding*); which is expected at this stage of the developmental cycle, especially with a ten-fold increase in EBs.

## Discussion

### How much influence is associated with increasing the MOI

There is a finite balance when infecting monolayers to accurately measure both host-cell and chlamydial transcriptional responses. This experiment used the universally standard MOI of 1, in addition to a ten-fold increase (MOI 10) and decrease (0.1), to directly observe what changes occur. One reason to increase the MOI is to examine early time points of infection when chlamydial transcripts are in low quantities as seen in (**Figure 2A**). In this experiment, when increasing the MOI to 10 at both time points, an increased capture rate of chlamydial transcripts was observed, confirming the suitability for early times (**Figure 2A-B**). However a challenge when working with higher MOIs is that some cells may have formed multiple inclusions which may skew host-cell responses beyond what may be seen in a real-world infection setting [52]. When looking at an MOI of 10 at 24 hours, we see over 60% of total captured transcripts from *Chlamydia*. Although this may not be representative of an *in vivo* infection, it is highly useful when focusing on chlamydial-based mechanisms. However, this does raise a question regarding a theoretical maximum proportion of chlamydial reads that can exist within a host cell during the developmental cycle, particularly at the later stages of infection. This was highlighted from (**Figure 1B**) at 24 hours, where we see a single replicate showing a staggering 74% of all transcripts associated with chlamydial expression. What was difficult to determine was if the increase in EBs had a corresponding influence in reducing the length of the developmental cycle due to possible synergistic interactions and shared resources. A future dual-RNA-seq study following the time course of [26], but replacing biovars for MOIs would be intriguing.

### What is an optimal MOI

We know from existing studies that different MOIs need to be used when examining different stages of the developmental cycle. For example, early time points generally require a higher MOI as limited transcription from *Chlamydia* occurs, resulting in low capture rates that can be difficult to interpret [19, 20]. During mid-stages, an MOI of 1 is often used to capture events based around growth and replication [53]. Towards the latter stages, almost all chlamydial genes are transcribed, making biological interpretations challenging [15]. Most studies examining a range of developmental stages use an MOI of 1 and this has generally been considered suitable. However, MOIs are generated from serial dilutions, so lower MOIs such as 1, may not actually have 1 EB per cell. Results from this experiment show a substantial increase in capture rates and transcription from an MOI of 1 to 10, suggesting that a slightly higher MOI may be optimal. There are however implications that need to be taken into consideration when using MOIs higher than 1. These include that EBs preferentially infect cells together rather than spread out evenly, which can result in all variations of the intended MOI. For example, a starting MOI of 5 will likely see an MOI range between 0-5 across a population of cells. As a result, the overall captured signal may be difficult to interpret, particularly with large MOIs. Furthermore, the length of the developmental cycle is generally shortened when many EBs are internalised due to an increased burden on the host cell.

### Combining depletion methods increases capture rate of chlamydial transcripts

By combining rRNA and polyA depletion methods, we clearly observe an increased capture rate of chlamydial transcripts. These additional transcripts do not appear to be from a small subset of genes dominating capture, but from a wide range expressed genes (**Figure 3**). However, host-based expression is affected, with expression of non-protein coding genes increasing (**Figure 4**); suggesting it may only be beneficial for future chlamydial-specific sequencing approaching to use both depletion methods.

### Experimental limitations

Unfortunately, this experimental design did not include any mock infected replicates from either time point, limiting some analyses. Future experiments would benefit from their inclusion, helping to separate general cell proliferation events from infection-relevant results.

A further limitation was not having the ability to determine if the timeframes of the developmental cycle are affected relative to increasing the MOI. The possibility of this occurring is quite likely, particularly as more EBs are internalised, resulting in more inclusions putting an increased burden on the host cell. Perhaps a future experiment could include a fluorescent tag that could be quantified as cells become lysed, thereby providing a measurement relative to the MOI and length of the developmental cycle. Alternatively, chemical-assisted methods can arrest at different cell cycle stages, ensuring all cells are synchronised at a specified cell cycle phase and removing this as a potential confounding factor.

### How dynamic is the chlamydial response

Differentially expressed and trended genes (**Figure 6**) identified transcriptional responses the host cell uses during an infection, which appears to be from a similar subset of key genes at both times. Genes are associated with immune related pathways, specifically inflammation; with increased expression as the MOI increases. As the concentration of EBs increases provoking this increased immune response, host cells will likely become overwhelmed if the numbers of EBs become too high. We hypothesise this could be an advantage for *Chlamydia* if a large proportion of host cell expression is focused towards immune responses, and if they already have a way of countering these, then other host processes may be easier to interfere with and possibly hijack. We anticipate this would most likely occur at higher MOIs where we have observed the most difference, particularly at 24 hours or latter stages of the developmental cycle.

## Conclusion

This work highlights how future bacterial-specific RNA-seq studies can increase sequence capture rates by combining rRNA and polyA depletion methods. This is particularly relevant for chlamydial-based expression studies when examining early time points, as low expression is generally observed. Three different MOIs highlighted that significantly more Chlamydial transcripts were captured at both time points when using an MOI of 10. Although useful for capturing Chlamydial-specific biology, the increased burden on host cells may not be representative of *in vivo* infections. Overall, these outcomes can help influence future NGS-based experimental designs to achieve more specific infection-related biological outcomes, particularly from *Chlamydia*-infected cells.

## Acknowledgements

This research was supported by UTS Faculty of Science Startup funding to GM. Sequencing was performed at the Genome Resource Centre, Institute for Genome Sciences, University of Maryland School of Medicine. Data was analysed on the ARCLab high-performance computing cluster at UTS, with files hosted using the SpaceShuttle facility at Intersect Australia

## Author Contributions

RH analysed, interpreted, and co-wrote the manuscript. MH performed the chlamydial infections and RNA-seq laboratory methods. WH assisted with interpretation of the data and contributed to the manuscript. GM conceived the experiments, obtained the funding, oversaw the sequencing, data analysis and interpretation, and co-wrote the manuscript.

## Supplementary files

**Supplementary File 1:**

Functional characterisation from top 25 highly expressed genes (host and chlamydial) that overlap all three MOI ratios.

